# RaftProt V2: understanding membrane microdomain function through lipid raft proteomes

**DOI:** 10.1101/286039

**Authors:** Ahmed Mohamed, Anup Shah, David Chen, Michelle M. Hill

## Abstract

Cellular membranes feature dynamic submicrometer-scale lateral membrane domainsvariously referred to as lipid rafts, membrane rafts or glycosphingolipid-enriched microdomains (GEM). In order to understand the molecular functions of lipid rafts, numerous studies have utilized various biochemical methods to isolate and examine the protein composition of membrane rafts. However, interpretation of individual raft proteomics studies are confounded by the limitations of isolation methods and the dynamic nature of rafts. Knowledge-based approaches can facilitate biological data interpretation by integrating experimental evidence from existing studies. To this end, we previously developed RaftProt (http://lipid-raft-database.di.uq.edu.au/), a searchable database of mammalian lipid raft-associated proteins. Despite being a valuable and highly used resource, improvements in search capabilities and visualisation were still needed. Here, we present RaftProt V2 (http://raftprot.org), an improved update of RaftProt, enabling interrogation and integration of datasets at the cell/tissue type and UniRef/Gene level. Besides the addition of new datasets and re-mapping of all entries to both UniProt and UniRef IDs, we have annotated the level of experimental evidence for each protein entry. The search engine now allows for multiple protein or experiment searches where correlations, interactions or overlaps can be investigated. The web-interface has been completely re-designed and offers new interactive tools for data and subset selection, correlation analysis and network visualization. Overall, RaftProt aims to advance our understanding of lipid raft function by revealing the proteomes and pathways that are associated with membrane microdomains in diverse tissue and conditions.

Database URL: http://raftprot.org

## INTRODUCTION

Biological membranes perform critical roles in compartmentalisation and signal transduction between the external environment and the cell, as well as between intracellular organelles. The existence of lateral membrane sub-compartments was initially suggested by the distinct glycolipid and glycosphingolipid-anchored proteins content on the apical and basal membranes of epithelial cells (1,2), leading to use of the term glycosphingolipid-enriched membranes (GEM). The ‘lipid raft’ hypothesis was developed in 1997 to explain the biophysical principles underlying the lateral membrane movements, and it proposed that the interaction between specific lipids drive the formation of functionally important lateral membrane domains (3). This hypothesis is supported by the biophysical properties of liquid-liquid phase separation, with cholesterol as one of the key lipids driving raft formation (4-6). The tight packing of raft lipids confers resistance to extraction by treatment with cold anionic detergent (Triton X-100) *in situ*, which, together with the demonstration of clustered glycosphigolipid-anchored proteins using immunofluorescence and immuno-electron microscopy, led to the interpretation of preferential localization of these proteins to lipid raft microdomains (7). Recent studies using super-resolution microscopy or single particle tracking indicate that lipid rafts are highly dynamic, transient nanometer-scaled membrane domains (8,9), present in numerous cellular organelles (10,11). Together, these studies support the potential role of lipid rafts in signal transduction by selective recruitment of transmembrane or peripheral membrane proteins with affinity for liquid ordered membranes (6,12-17).

An important step towards understanding the molecular mechanisms and effectors of lipid raft function is the characterisation of its proteomic compositions in various cell types and conditions. Given the dynamic nature and nanometer scale of lipid rafts, isolation of individual or pure lipid raft microdomains for proteomics analysis is beyond the capabilities of current techniques. On the other hand, numerous studies have reported the proteomic analysis of bulk lipid rafts prepared from diverse cell and tissue types, and these public datasets are a useful resource for knowledge-based data mining and integration. To facilitate this, we developed and reported RaftProt, a searchable database for mammalian lipid raft proteomics data (18). We used RaftProt data to provide independent experimental support for lipid raft localization of IQGAP1, its involvement in cancer metastasis (19). Since publication, RaftProt database has been used to compare with new lipid raft proteomics data from bovine lens (20), ovarian cancer cells (21) and mouse brain (22), as well as HeLa cell surface proteome data (23) and in a computational study of single-pass transmembrane proteins (24).

To further the utility of RaftProt, we implemented several new features. Firstly, we mapped each UniProt ID to their UniRef and GENE ID, to allow cross-species and gene-level interrogation., Furthermore, we re-designed the web-interface to provide interactive tools for dataset selection, analysis and visualisation. A new experimental evidence level annotation has been developed and implemented, providing a simple summary score for each protein entry. In addition, the database has been relocated to a simpler domain name (http://raftprot.org).

## DATABASE DESCRIPTION

RaftProt V2 has been completely re-developed, and the web address updated from http://lipid-raft-atabase.di.uq.edu.au/ to http://raftprot.org. We describe below improvements in the data content, visualization and usability.

### Database architecture

RaftProt V2 is implemented with a set of RESTful Web APIs, as shown in **Figure 1**. When users use RaftProt directly via a web browser, all queries are issued to the database via this Web API. Furthermore, this Web API is open to public access, hence users can write their own programs (in the programming language of their choice) to retrieve the most up-to-date data from our database, and perform their own customised analysis. The Web API can return data in different formats such as CSV, XML, or JSON. RaftProt V2 web site contains detailed API documentation, along with helpful code examples. In addition, users can browse the API and test different queries to help with their implementation.

**Figure 1.**
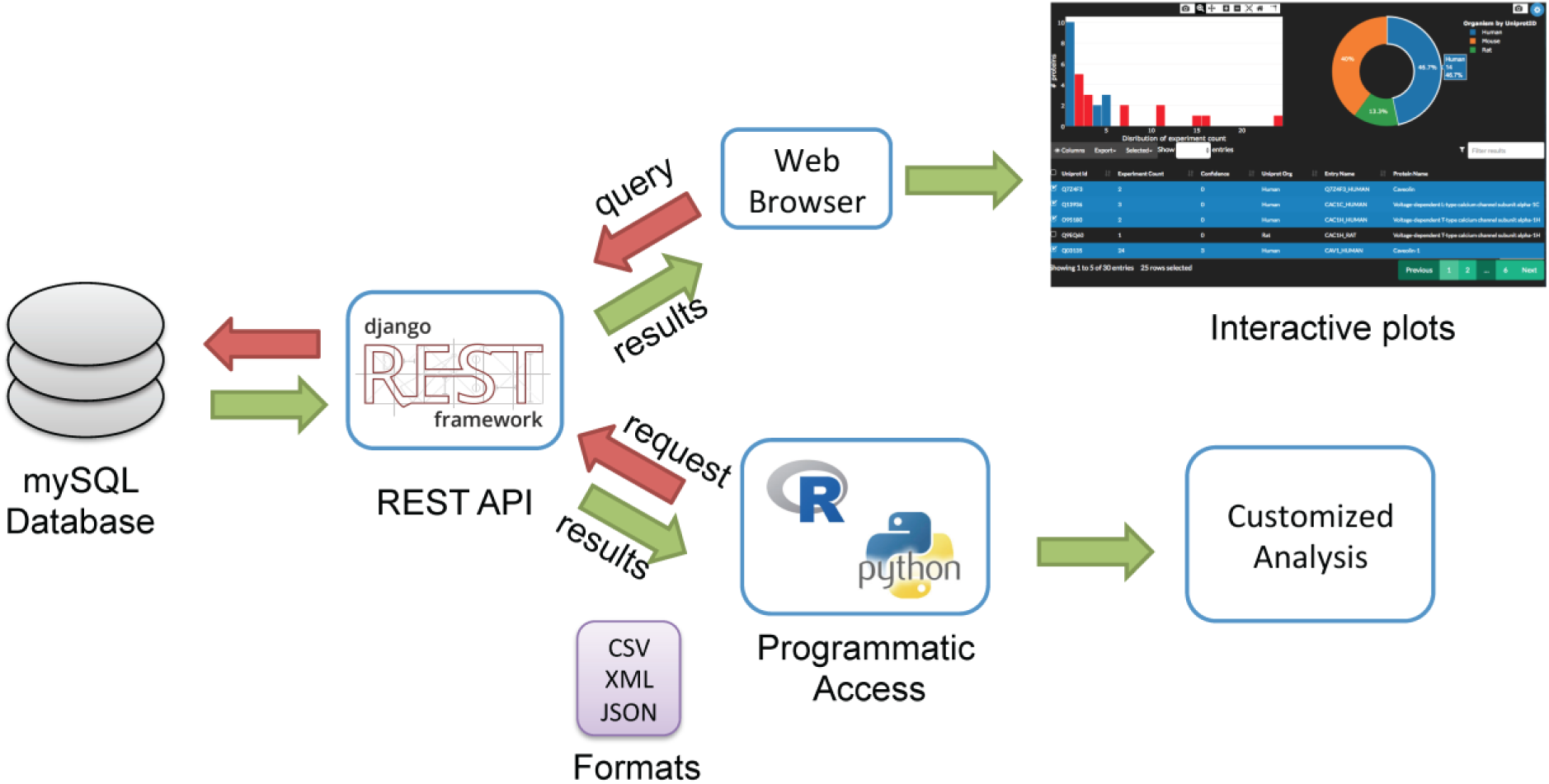
RaftProt V2 Database architecture.

### Data collection and curation

New lipid raft proteomics datasets from July 2014 to June 2017 were identified through PubMed using the terms ‘lipid raft’ or ‘microdomains’ with ‘proteomics’ or ‘proteome’. Data were downloaded from online supplemental information of articles, or requested from corresponding authors. RaftProt V2 currently includes 98 studies and 163 experiments from 6 species. Human lipid raft proteome is the most studied with more than 50% of experiments, followed by mouse and rat.

In addition to supplementing the database with new studies, we also updated the annotation for previously collected data. All protein IDs were remapped to the latest version of Uniprot database. Obsolete Uniprot IDs were updated where possible or removed. We flagged IDs as “mismatching” whenever they belonged to different species than the reporting experiment. Experiments with more than 50% mismatching IDs were also flagged. Flagged proteins and experiments were excluded from the subsequent evidence levels scoring. Finally, we added UniRef IDs for all mapped proteins, which clusters proteins according to their sequences, thereby allowing users to explore closely related proteins in the lipid rafts.

### Experimental evidence level annotation

Due to its simplicity, detergent-resistant membrane (DRM) extraction method has been widely used to infer the presence of proteins in lipid rafts (7). While most of the experiments in RaftProt use DRM method for extraction, additional methods have also been reported, for example, detergent-free methods relying solely on the buoyancy of membranes in a sucrose gradient (25), and affinity-based methods (26). In addition, sensitivity to cholesterol extraction by methyl-β-cyclodextrin has been proposed as the gold standard for defining *bona fide* lipid raft resident proteins in cell culture-based studies (8). The biochemical extraction procedures have been captured for each experiment. In the previous version of RaftProt, a searchable list of high confidence lipid raft proteins was generated, being the subset of proteins reported using more than one extraction method OR reported to be sensitive to methyl-β-cyclodextrin treatment (18). To improve the flexibility of database searching, we have now implemented an experimental evidence level rating system. Each entry is annotated with upto 3 stars, where 1 star denotes identification by more than one biochemical extraction method, 2 stars denote sensitivity to methyl-β-cyclodextrin, and 3 stars denote compliance with both criteria. Zero star indicates that the protein has only been reported using a single biochemical extraction method (usually DRM) and has not been reported to be sensitive to methyl-β-cyclodextrin (either due to lack of such experiment, or actually being insensitive).

**Figure 2a** shows species-specific number of UniProt IDs for each experimental evidence level from 0 to 3. More than 30% of the reported human membrane raft proteins scored 1 star or more. Surprisingly, more than 80% of reported mouse, rat and bovine membrane raft proteins scored 0, possibly reflecting a large proportion of species-specific UniProt IDs being reported in only a small number of studies, and the exclusive use of DRM method.

**Figure 2.**
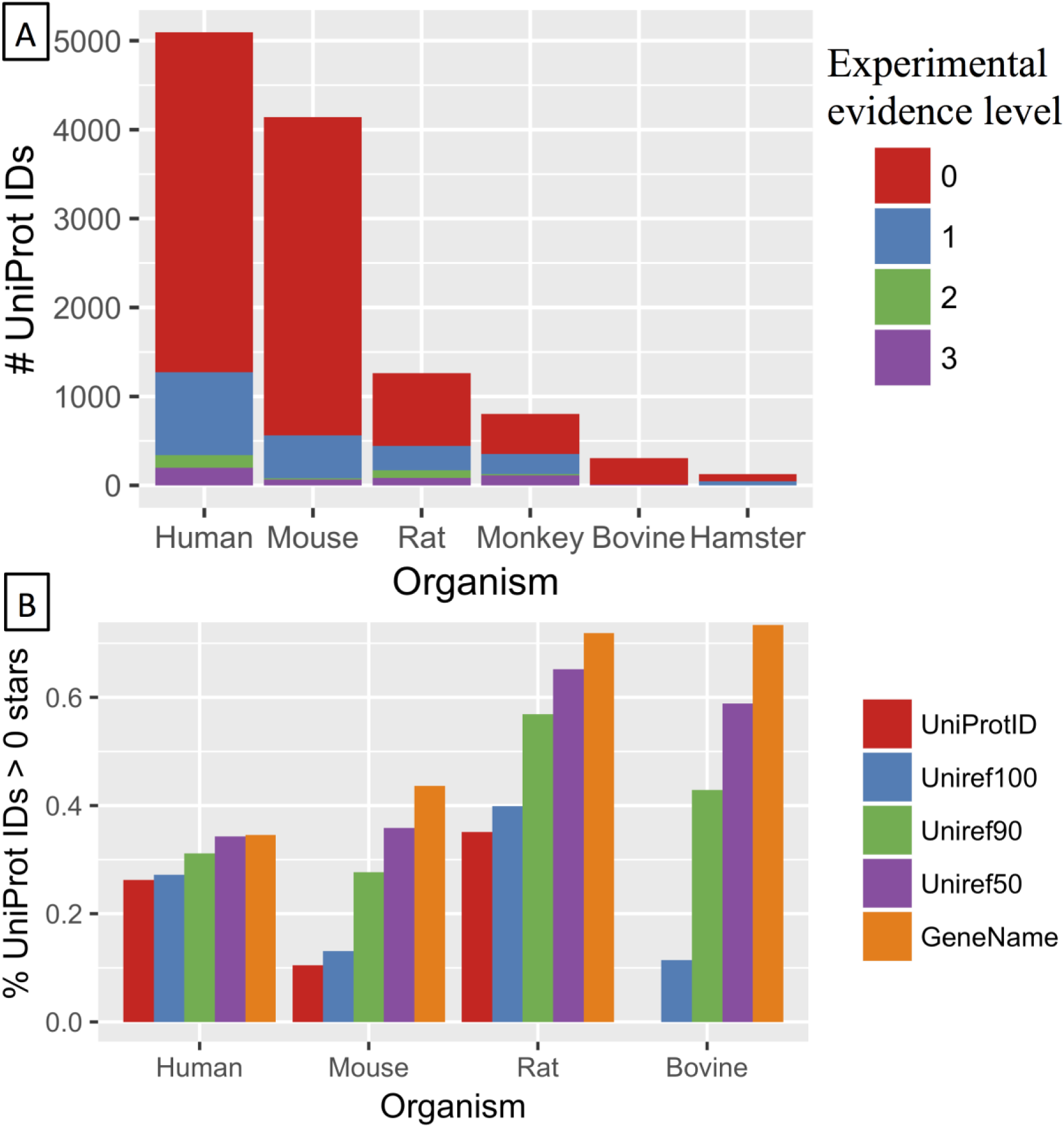
Distribution of lipid raft protein experimental evidence levels by organism. **A** Number of UniProt IDs with each experimental evidence level. **B** Percentage of UniPort IDs with more than zero stars for each calculated experimental level.

### Cross-species analysis

For each protein entry, users can examine closely related proteins, i.e. sharing the same UniRef ID or gene name, using the Cross-species info tab. Users can further evaluate experiments reporting these proteins by directly clicking on the displayed search icons. To collate experimental evidence levels for proteins identified in several species, each UniProt entry was further annotated with experimental evidence levels based on UniRef IDs and gene name. The aggregation of experimental evidence across different species improves the scores greatly, especially in species with limited number of experiments available in the database (**Figure 2b**).

## DATABASE USAGE

The user interface has been completely re-designed to improve data analytics and visualisation. All search results are now displayed as interactive plots using JavaScript libraries. The interactive plots and tables allow users to subset and select from search results using different parameters such as experimental evidence level, biochemical extraction or quantitative proteomics methods. Output tables and plots can be exported, and protein lists can be saved for use in later searches.

### Searching

RaftProt V2 allows users to query the database using any of the four entities, proteins, experiments, studies or tissues, using two different search types, single and multiple (**Figure 3**). Single protein searches are performed using the “Quick search” interface. Outputs from RaftProt single protein searches include statistics of experiments in which the selected protein was identified, where experiment subsetting can be performed by species or quantitation method, as well as the list of proteins co-identified with the query protein displayed in a table and interactive frequency graph.

**Figure 3.**
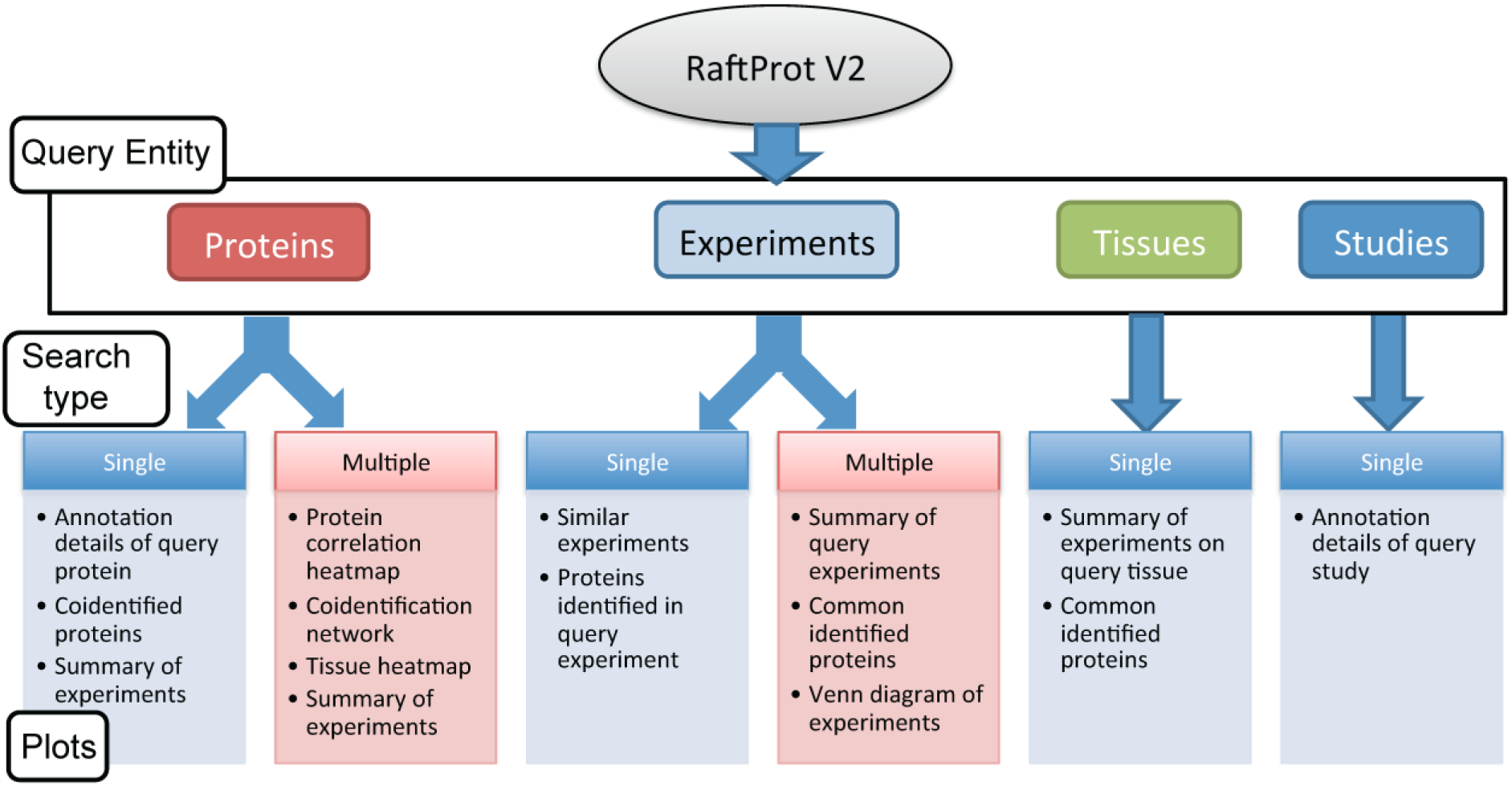
RaftProt V2 search options and data output.

The “Advanced search” interface allows users to query a list of proteins, such as protein family or results of a new experiment, to investigate their relationship in membrane raft. In addition to providing a summary of experiments which report the list of proteins, RaftProt V2 offers three types of correlation tools for multi-protein searches; pairwise correlation, network visualisation and protein-tissue heatmap. Use of these visualisation tools is exemplified in **Figure 4** with caveolin and cavin family proteins. Searching was performed using a list consisting of CAV1, CAV2, CAV3, CAVIN1, CAVIN2, CAVIN3 and MURC. The heat map in **Figure 4A** shows that CAV1, CAV2, CAVIN1, CAVIN2 and CAVIN3 are frequently co-identified in many membrane raft datasets, but CAV3 and MURC are less commonly co-identified with the other family members. These results are expected since CAV3 and MURC are both predominantly expressed in muscle, while the other family members have broad tissue expression (27). Extending beyond pairwise comparison, the network of the co-identification of these proteins are shown in **Figure 4B** as a static image. Using the interface, hovering over the nodes will displays the accession number and highlights the table row for the protein entry. The thickness of the edges correlates with the number of datasets that identified the connected nodes. **Figure 4B** shows two major hubs, one less densely connected than the other, representing the muscle and non-muscle caveolin-cavin complexes, respectively. Finally, the protein-tissue heatmap in **Figure 4C** further confirm the muscle-specific expression of CAV3 and MURC, and also reveal interesting expression patterns for the other family members.

**Figure 4.**
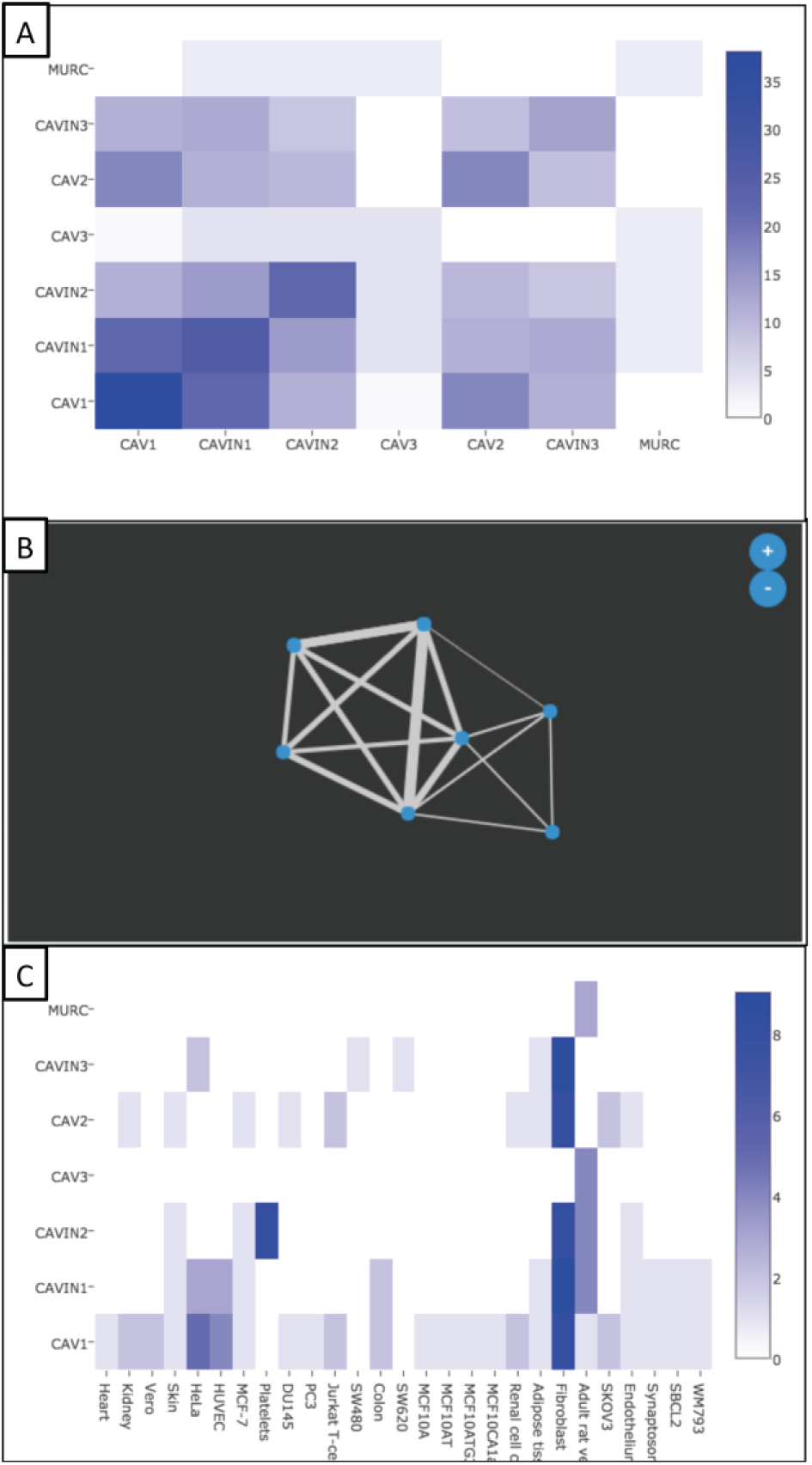
Example correlation analysis outputs using the list CAV1, CAV2, CAV3, CAVIN1, CAVIN2, CAVIN3, MURC. **A**. Heatmap showing pairwise co-identification of proteins. **B**. Network visualization for co-identification of proteins. **C**. Protein-tissue heatmap showing co-identification in different tissues.

### Browsing

Users can also browse experiments or studies according to species or tissue/cell type. The latter is visualised by an ontology tree which was developed for RaftProt (18), and updated with new tissue types. Experiments annotated with the selected tissue/cell type are displayed in a table with interactive donut graphs showing experimental organism and quantitation method. The user can easily select the segment of choice and subset those experiments using the Selected → Search selected option. Common proteins of the selected experiments are displayed in frequency a graph under the ‘Commonly Identified Proteins’ tab, while the ‘Experiments Venn’ tab visualises the overlap in the lipid raft proteome between the selected experiments. There is a limit of 10 experiments for the Venn diagram visualisation.

## DATABASE DOWNLOAD

All data included in RaftProt V2 can be downloaded from the website. Users can choose to either download the whole database, or a subset based on the organism and/or experimental evidence levels. Data can also be downloaded in several file formats including CSV and XML.

## CONCLUSIONS AND FUTURE DIRECTIONS

RaftProt V2 is a searchable database for the integrated analysis of mammalian lipid raft proteomics datasets. Future expansion of RaftProt may include proteomics datasets uploaded into data repositories but not linked to a publication, non-mammalian lipid raft proteomic studies or lipidomics studies, as well as additional analytical and visualisation tools.

## ACKNOWLEDGEMENT

We thank Mr Anthony Thyssen (Griffith University) for assistance with server setup, and the authors who have kindly provided proteomics data summaries upon request.

